# Time-dependent-asymmetric-linear-parsimonious ancestral state reconstruction

**DOI:** 10.1101/089805

**Authors:** Gilles Didier

**Affiliations:** Aix-Marseille Université, CNRS, Centrale Marseille, I2M, UMR 7373, 13453 Marseille, FRANCE

**Keywords:** maximum parsimony, ancestral state reconstruction, quantitative characters, parametric methods

## Abstract

The time-dependent-asymmetric-linear parsimony is an ancestral state reconstruction method which extends the standard linear parsimony (a.k.a. Wagner parsimony) approach by taking into account both branch lengths and asymmetric evolutionary costs for reconstructing quantitative characters (asymmetric costs amount to assuming an evolutionary trend toward the direction with the lowest cost).

A formal study of the influence of the asymmetry parameter shows that the time-dependent-asymmetric-linear parsimony infers states which are all taken among the known states, except for some degenerate cases corresponding to special values of the asymmetry parameter. This remarkable property holds in particular for the Wagner parsimony.

This study leads to a polynomial algorithm which determines, and provides a compact representation of, the parametric reconstruction of a phylogenetic tree, that is for all the unknown nodes, the set of all the possible reconstructed states associated to the asymmetry parameters leading to them. The time-dependent-asymmetric-linear parsimony is finally illustrated with the parametric reconstruction of the body size of cetaceans.

## 1 Introduction

Testing hypotheses about evolutionary mechanisms like environment influence, homoplasy etc. calls for information not only about the extant organisms but also about the ancestral ones, which is mostly inaccessible, with a few exceptions when related fossils can be found. More generally, how to study the evolution since we cannot observe the process ongoing over a significant time scale, but only see its result? A common way to cope with this issue is to infer the unknown, and essentially unknowable, information about the ancestral organisms from that observed on the extant taxa, by using the phylogenetic relationships between these last ones. Such an inference is sometimes called *character mapping* or more often, *ancestral state reconstruction* [12, 7, 2]. Note that the problem of determining the phylogenetic relationships between extant taxa is generally treated independently of, and prior to, the ancestral reconstruction. In short, ancestral state reconstruction approaches generally assume that both the phylogeny and the character states of extant taxa are given and aim to infer the ancestral states from these data.

Ancestral reconstruction is a challenging and important question in evolutionary biology. As such, it motivated the development of several approaches [5, 9, 8, 6, 13]. They all rely on two general points of view which are quite different. Namely, ancestral reconstruction methods are based either on the parsimony principle or on stochastic models of character evolution. Note that reconstruction approaches differs not only in the principle underlying them, but also in the nature of the states that they aim to reconstruct. One does not use the same methods for reconstructing continuous/quantitative features (e.g. size, weight, cranial volume…) or discrete characters which are here the characters taking only a finite number of values like the number of fingers, the presence or absence of a given feature etc. In [12], authors differentiate discrete characters according to the way in which their possible values can follow one another during evolution. For instance, *linear ordered characters* have ordered values and are such that evolving from a value to another requires to pass through all the intermediate values.

The present work focuses on the parsimonious method for reconstructing continuous character in which the cost of an evolution is the absolute difference between its ending and starting states. This method is referred to as the Wagner or the linear parsimony [4, 11, 1]. Following [1], we consider an asymmetric version of this method, allowing the cost of an increasing evolution to be different from that of a decreasing evolution of the same amount. Moreover, we allow the cost of a character variation during an interval of time, to depend on the length of this interval (in practice, lengths of intervals of times are branch lengths). The reconstruction method such obtained is called *time-dependent-asymmetric-linear parsimony* (*TDALP*).

Our starting point is a detailed study of the influence of asymmetry parameter on the reconstructed states. This study is based on the characterization of the function associating a state *x* and an asymmetry parameter *γ*, with the smallest cost, under *γ*, of a reconstruction forced to infer *x* at the root of the tree [1]. The properties of this function allow us to prove that, whatever the asymmetry parameter, there always exists a reconstruction in which all the inferred states are taken among the known states (generally those of the tips, but we make no assumption on this point). Reconstructions containing states which are not among the known ones are degenerates cases which only occurs for a finite number of special values of the asymmetry parameter. This strong property, which is much stronger than saying that the reconstructed states lies in the range of the known ones, makes the approach totally relevant to deal with discrete characters, in particular with linear ordered ones. To our knowledge, the TDALP is the only reconstruction approach which can be applied to both discrete and continuous characters. For instance, reconstruction methods based on stochastic models handling continuous character use Brownian motion to model their evolution, and are very different to those dealing with discrete characters which use Markov models for the same purpose.

The results obtained about the influence of the asymmetry parameter are next used for designing an algorithm which determines, for all unknown nodes of the tree, the different states reconstructed by the TDALP according to the asymmetry parameter. For an unknown node, the set of all the possible reconstructed states with the corresponding asymmetry parameters, is called its *parametric reconstruction*. The algorithmic complexity of the algorithm is polynomial (quartic with the size of tree) both in time and memory space. The TDALP can be applied to a wide variety of biological datasets for reconstructing characters of any nature whatsoever (see above).

I developed a software implementing the approach presented here. This software takes as inputs a phylogenetic tree with or without branch lengths, and a file containing the known character states (in standard “.csv” table format) and outputs the parametric reconstruction of all the unknown nodes in several formats, notably in graphical ones as displayed in Figure. 2 and 3. Its source code, written in C language, is freely available at https://github.com/gilles-didier/TDALP.

The rest of the paper is organized as follows. We introduce the notations, a formal presentation of the ancestral reconstruction problem and the first definitions in Section 2. The detailed study of the influence of the asymmetry parameter is carried out in Section 3. It leads to the definition of the parametric reconstruction of a character at a node, which is formally defined at the beginning of Section 4. This section continues with the presentation of an algorithm computing the parametric reconstruction of all the unknown nodes of a given tree and the study of its complexity. We conclude with Section 5 by illustrating our approach with the reconstruction of the body size of cetaceans.

## 2 Definitions and notations

The cardinal of a finite set *S* is noted |*S*|.

Let *T* be a rooted tree which is not required to be binary. As it should lead to no confusion, we still write *T* for its set of nodes. For all nodes *n* ϵ *T*, we put

- *C*_*n*_ for the set of child nodes of *n*,
- *τ_η_* for the length of the branch ending at *n*,
- *a*_*n*_ for the direct ancestor of *n*,
- *T*_*n*_ for the subtree of *T* rooted at *n*.

Before introducing the reconstruction problem, let us start by considering a subset 𝒦 of of nodes of *T*, a map *ϑ* from 𝒦 to the set of real numbers R. The map *ϑ* will be referred to as the *initial function* and the nodes of 𝒦 are said *known*. In the standard ancestral state reconstruction problem, 𝒦 is exactly the set of tips of *T* but we make here no assumption on 𝒦. In plain English, any node of the tree can be known or not. For all nodes *n* of *T*, we put 𝒦_*n*_ for the subset of known nodes of the subtree *T*_*n*_, i.e. 𝒦_*n*_ = 𝒦 ∩ *T*_*n*_. The values of {*ϑ* (*k*) | *k* ϵ 𝒦} are the *known states* of *T*. For all nodes *n*, we put *ϑ* (𝒦_*n*_) for the set {*ϑ* (*k*) | *k* ϵ 𝒦_*n*_}.

For technical reasons, we will consider two special nodes *o* and *p* not belonging to *T*, for which, by convention, we set *ϑ* (*o*) = -∞ and *ϑ* (*p*) = +∞.

The *ϑ-assignments* of *T* are the maps *ξ* from *T* to the set of real numbers which extend *ϑ* (i.e. such that *ξ*(*n*) = *ϑ*(*n*) for all nodes *n* ϵ 𝒦). The reconstruction problem of *T* with regard to *ϑ* consists in finding the most relevant (in some sense) *ϑ*-assignment of *T*.

### 2.1 Parsimonious reconstruction

In the parsimony framework, the relevance of an assignment is expressed in terms of cost. More specifically, in the time-dependent-asymmetric-linear parsimony (*TDALP*) case, an ancestor/child transition from value *x* at node *n* to value *y* at its child m is associated with the cost:

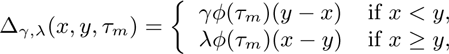

where *λ* and *γ* are two positive real numbers and *ϕ* is a function from ℝ_>0_ to ℝ_>0_. In what follows, we make the assumption that *τ_n_* > 0 for all nodes *n* ϵ *T*. In other words, we forbid null branch lengths (but we allow polytomies). Remark that the generalized linear parsimony as defined in [1], which generalizes the Wagner parsimony [4], corresponds to the case where *ϕ* is constant with *ϕ*(*τ*) = 1 for all *τ*. In an evolutionary context, it makes sense to choose a decreasing function *ϕ* (roughly speaking, evolving takes times), but any positive function can be used. In Section 5, we set 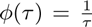 for reconstructing the body size of cetaceans. The special case where *γ* = *λ* = 1 and *τ*(*τ*) = 1 for all *τ*, corresponds to the Wagner parsimony. Remark that steadiness is always cost-free, i.e. Δ_*γ,λ*_(*x,x,τ*) = 0 whatever *x, τ, γ, λ* and *ϕ*.

The cost Δ_*γ,λ*_(*ξ*) of an assignment *ξ* of *T* is then the sum of all the costs of its ancestor/child transitions:

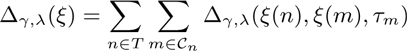

Finding the most time-dependent-asymmetric-linear parsimonious reconstruction on *T* with regard to *ϑ* consists in determining a *ϑ*-assignment ξ of *T* with a minimal cost Δ*_γ,λ_*(*ξ*).

Let us first remark that multiplying both *λ* and *γ* by a positive constant factor does not change the relative order of the assignments costs. Since *λ* > 0, we can divide both parameters by *λ* without changing which assignments are the most parsimonious. From now on, we assume without loss of generality that *λ* = 1. The cost of an assignment only depends on the parameter *γ*, and is called *γ-cost*:

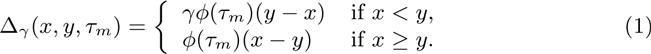

Below, *γ* will be referred to as the *asymmetry parameter.* Intuitively, reconstructing with *γ* < 1 (resp. with *γ* =1, with *γ* > 1) corresponds to the assumption that the character evolves with a positive trend (resp. without trend, with a negative trend).

A *γ-parsimonious reconstruction* of *T* with regard to an initial function *ϑ* is an *ϑ*-assignment with a minimum *γ*-cost.

### 2.2 Cost and stem cost functions

Let us borrow and adapt some definitions of[1]. For all nodes *n* ϵ *T*, the *(subtree) costfunction f*_*n*_ maps a pair (*γ, x*) ϵ ℝ_>0_ × ℝ to the smallest *γ*-cost which can be achieved by an assignment ξ of the subtree *T*_*n*_ such that *ξ*(*n*) = *x*. An assignment *ξ* of *T*_*n*_ satisfying *ξ*(*n*) = *x* will be said *x-rooted.* By construction, *f*_*n*_ (*γ x*) is thus the *γ*-cost of a *x*-rooted *γ*-parsimonious assignment of *T*_*n*_. By convention, if there exists no *x*-rooted assignment of *T*_*n*_ (typically when *n* ϵ 𝒦 with *ϑ*(*n*) ≠ *x*) then we set *f*_*n*_(*γ, x*) = +∞.

Claim 1. *For all γ* > 0 *and all real values x, we have that*

1. *if n is an unknown leaf then f*_*n*_(*γ, x*) = 0;
2. *if n is a known leaf then*

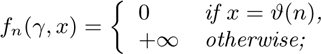
3. *if n is an unknown internal node then*

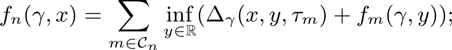
4. *if n is a known internal node then*

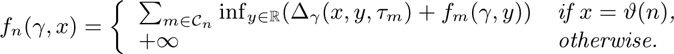

Items 3 and 4 of Claim 1 lead us to introduce an additional notation. For all non-root nodes *m* of *T*, we define the *stem cost function* 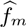 as

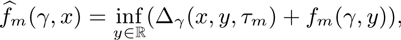

which can be understood as the smallest *γ*-cost which can be achieved by an assignment of the tree only made of the direct ancestor of *m* and of the subtree *T*_*m*_ which associates the direct ancestor of *m*, with the state *x*. Equation of Item 3 in Claim 1 becomes

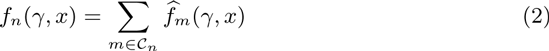

For all non-root nodes *m* of *T*, the tree made of the direct ancestor of *m* and of the subtree *T*_*m*_ will be referred to as the *stem-subtree* of *m* and noted 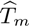.

Proving properties of *f*_*n*_ and 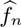 shall follow a same general induction scheme which will be referred to as the *standard induction scheme*. It comes from Lemma 1 of [1] and stands in the three following steps:

*Step 1*: establish the property for the base cases of 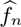 and *f*_*n*_;

*Step 2*: prove that if the property holds for *f*_*n*_ then the same property holds for 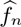;

*Step 3*: prove that if the property holds for all 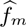 where *m* is a child of *n* then it holds for *f*_*n*_.

Steps 1 and 3 are generally plain. Both maps *f*_*n*_ and 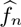 have very simple forms for the base cases (i.e. the leaves). In most cases, Step 3 follows straightforwardly from Equation 2. The main point of the proofs actually stands in Step 2.

## 3 Parametric analysis

We shall study the influence of the parameter *γ* on the states inferred in the *γ* parsimonious reconstruction. To this end, we start by studying the nature of cost and stem cost functions.

### Theorem 1.

*Let n be an unknown node of T. The maps f*_*n*_ *and* 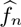 *are piecewise-linear and continuous and, for all γ* > 0, *the maps x → f*_*n*_(*γ,x*) *and* 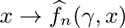 *are both convex*.

*More precisely, if all the nodes of T*_*n*_ *are unknown then f*_*n*_(*γ,x*) = 0 *for all γ* > 0 *and all x. Otherwise, there exist:*

- *an integer u*_*n*_ *and a strictly increasing positive real sequence* 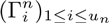,
- *an integer sequence* 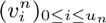 *and for, all* 0 ≤ *i* ≤ *u*_*n*_, *a sequence* 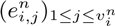 *of known nodes of T*_*n*_ (*i.e. of 𝒦*_*n*_) *verifying* 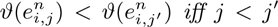*; by convention we set* 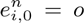 *and* 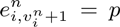 *(we recall that ϑ*(*o*) = –∞ *and, ϑ*(*p*) = +∞),
- *two non-negative real bi-sequences* 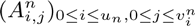*· and* 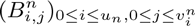,
- *two real bi-sequences* 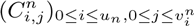 *and* 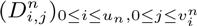

*such that, by setting* 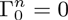 *and* 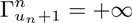 *and for all* 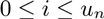, all 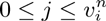, *all* 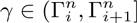 *and all* 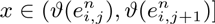, *we have*

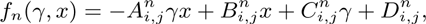

*all the coefficients being such that f*_*n*_ *is continuous. Moreover, the sequence* 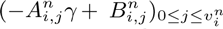 *(i.e. the x-coefficients of f*_*n*_*) is increasing with* 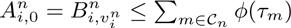 *and* 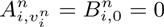.

*In the same way, if all the nodes of T*_*n*_ *are unknown then* 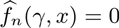 *for all γ > 0 and all x. Otherwise there exist* 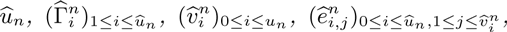 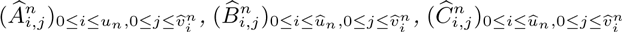 and 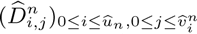 *verifying the same properties as their f*_*n*_*-counterparts except that we have* 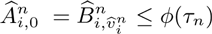, *and* 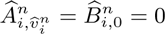 *and such that, for all* 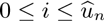, *all* 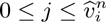, *all* 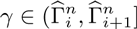 *and all* 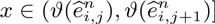, *we have, we have*

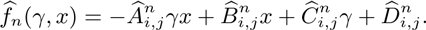

*Proof.* We follow the standard induction scheme and start with Step 1. The base cases of *f*_*n*_, i.e. when *n* is a leaf, are given by Items 1 and 2 of Claim 1. Let us check the base cases of 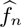 = 0. If *n* is an unknown leaf then we have 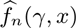 for all pairs *γ* > 0 and all *x*. If *n* is a known leaf then we have:

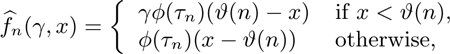

for all *γ* > 0 and all *x*. In all cases, 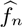 is piecewise linear and continuous. Moreover, we remark that, for *x* small enough (resp. large enough), the *x*-coefficients of *f*_*n*_ and 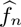 are either 0 or –*γϕ*(*τ*_*η*_) (resp. either 0 or *ϕ*(*τ*_*η*_)), according to whether *n* is unknown or not.

Let us proceed to Step 2 and assume that *n* is an internal node and that the theorem holds for *f*_*n*_. If *n* is known then we have:

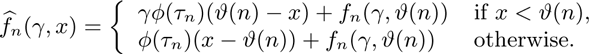

Since *f*_*n*_ is piecewise linear and continuous, the map 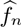 is well piecewise linear and continuous with 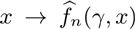 convex. The same remark as just above about the *x*-coefficient of 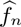 holds again.

Let now assume that *n* is unknown internal node. We still assume that the theorem holds for *f*_*n*_ and we assume in addition that *T*_*n*_ contains at least one known node (the other case being straightforward). Let us consider an index 0 ≤ *i* < *u*_*n*_ and a real number 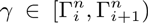. With the induction hypothesis, the map *y* → *f*_*n*_(*γ, y*) is convex, continuous and piecewise linear. So is the map defined for any real value *x* and all *y* ≥ *x* by

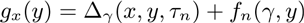

For all real values *x* and *y* ≥ *x*, we have that *g*_*x*_(*y*) = *γϕ*(*τ*_*n*_)(*y* – *x*) + *f*_*n*_(*γ, y*), which, under the notations of the theorem, can be written as:

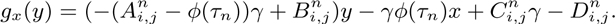

From the induction hypothesis, the sequence 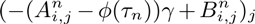 increases with *j*. Let 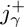 be the smallest index such that 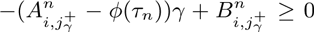 (i.e. that corresponding to the first interval on which *g*_*x*_ does not decrease with *y* ≥ *x*). Since, from our induction hypothesis, we have that 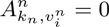, such an integer always exists and we get that

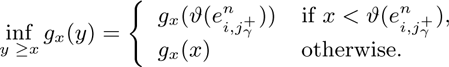

Let us remark that

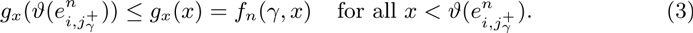

Symmetrically, for all real values *x* and *y* ≤ *x*, we have that *g*_*x*_(*y*) = *ϕ*(*τ*_*n*_)(*x* – *y*) + *f*_*n*_(*γ, y*), i.e.

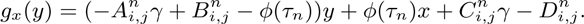

Let us define 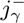 as the greatest integer smaller or equal to 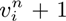 and such that 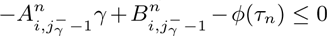 (i.e. that corresponding to the last interval on which *g*_*x*_ does decrease with *y* ≤ *x*). Since, from our induction hypothesis, we have that 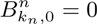, such an integer always exists. We have that

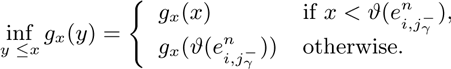

Let us remark that

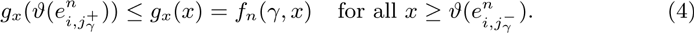

Since both *γ* and *ϕ*(*τ*_*n*_) are positive, we have that

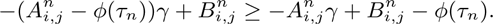

It follows that 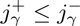 therefore 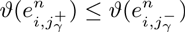.

Moreover, since

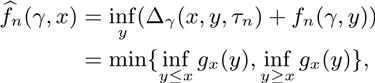

the inequalities 3 and 4 imply that

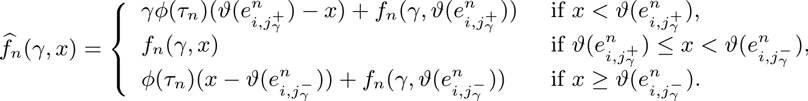

It straightforwardly follows that, for all *γ*, the map 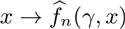 is piecewise linear and continuous. From the induction assumption, *x* → *f*_*n*_(*γ,x*) is convex, in particular between 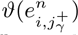 and 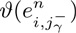. Still from the induction assumption, we have that 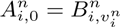 and 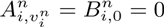. Two possibilities arise:

- If 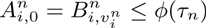 then both 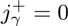 and 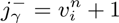. We have 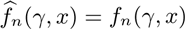. From the induction hypothesis, the function 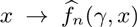 is well convex and such that 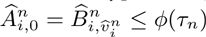 and 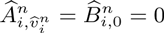.
- If 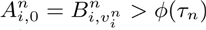, then we have both 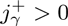 and 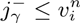. The definition of 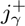 and of 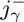 ensures that 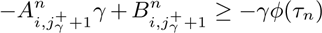 and 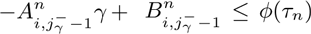, which implies that the map 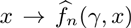 is still convex, in this case with 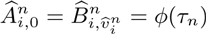 and 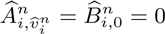.

Let us show that 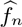 is piecewise linear with regard to *γ*. To do so, we put Φ_1_,…, Φ_*p*_ for the elements of

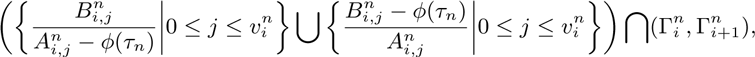

indexed in increasing order. By construction, the indexes 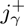 and 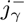 are both constant over all the sub-intervals 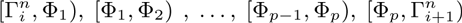. Since from our induction hypothesis *γ* → *f*_*n*_(*γ,x*) is linear over 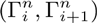 and for all values *x*, the map 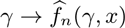 is linear over all the sub-intervals above. It follows that the map 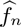 is piecewise linear and its continuity with regard to *γ* is straightforward to verify at all bounds Φ_*k*_ for 1 ≤ *k* ≤ *p*.

Step 3 is the last one remaining. Let *n* be an unknown internal node and let us assume that the theorem holds for all children *m* of *n*. In particular, the maps 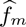 are piecewise linear, continuous and convex with regard to *x*. As sum of these maps (Equation 2), the map *f*_*n*_ satisfies itself these properties. Moreover, since for all children *m* of 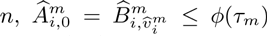 and 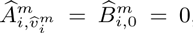, we have that 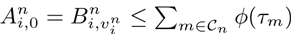 and 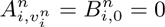.

Figure 1 shows an example of cost function and its graphical representation.

For all functions *f*_*n*_ (resp. 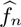) and all 0 < *i* < *u*_*n*_ (resp. all 0 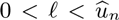), the bounds 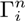 (resp. 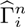) will be referred to as *asymmetry-bounds* of *f*_*n*_ (resp. of 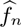) and, for all 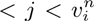 (resp. all 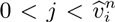), the bounds 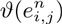 (resp. 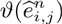) will be referred to as *state-bounds* of *f*_*n*_ (resp. of 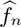).

**Remark 1**. *In the generalized parsimony scheme of [1] (i.e. when ϕ(τ) = 1 for all τ), both* 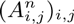 *and* 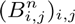 *are integer bi-sequences and the sequence* 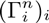 *is rational*.

Since *ϕ*(*τ*_*k*_) is assumed positive for all nodes *k* ∈ *T*, the following remark is straight-forward to prove by induction.

**Remark 2**. *If n is an unknown node such that* 𝒦_*n*_ ≠ ∅, *there exists no pair* (*i, j*) ∈ 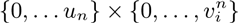 *such that* 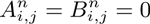.

The proof of the following corollary is essentially contained in that of Theorem 1.

**Corollary 1**. *Let γ be a positive real number, n a node of T, m a child of n and, under the notations of Theorem 1, i be the greatest index such that* 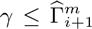. *If a γ-parsimonious reconstruction assigns the value x to n then it assigns to all unknown children m of n*,

1. *a value between* 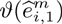 *and* 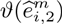 *(both included) if* 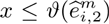 *and* 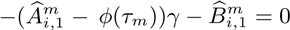,
2. 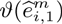 *if* 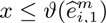 *and* 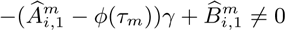,
3. *a value between* 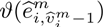 *and* 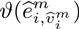 *(both included) if* 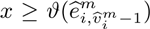 *and* 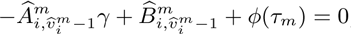,
4. 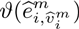 *if* 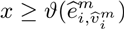 *and* 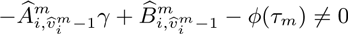.
5. *x otherwise*.

*Cases 1 (resp. Case 3) occurs if and only if* 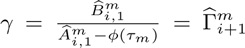 (*resp*. 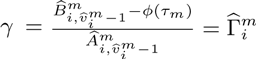). *In particular, if we have both* 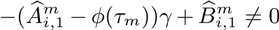 *and* 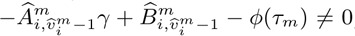, *which is always true if* 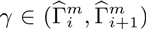, *then if a γ-parsimonious reconstruction assigns the value x to n then it assigns to all unknown children m of n*,

- 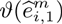 *if* 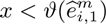,
- *x if* 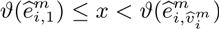,
- 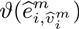 *if* 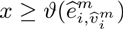

(*this point was already stated in [1]*).

### Theorem 2.

*For all asymmetry parameters γ, there exists a γ-parsimonious reconstruction in which all the reconstructed states belong to ϑ*(𝒦). *Moreover, if a reconstruction is both γ- and γ′-parsimonious for two positive real numbers γ ≠ γ′ then its inferred states all belong to ϑ*(𝒦).

*Proof.* Let us start by proving that for all *γ* > 0, there exists a *γ*-parsimonious reconstruction which assigns a known value to the root. This is plain if the root *r* is known. If *r* is unknown then, under the notations of Theorem 1 and for all *γ* > 0, the map *x* → *f*_*r*_ (*γ, x*) is piecewise linear, continuous and convex with

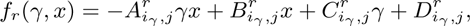

for all 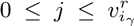 and all 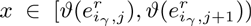, where *i*_*γ*_ is such that 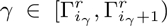.

Let *j*_*γ*_ be the smallest index such that 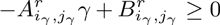, which always exists (Theorem 1). Since the sequence 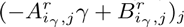 is increasing, we have that

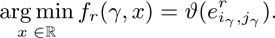

Moreover, there exists a real 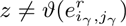 such that arg min 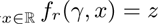 if and only if 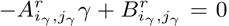. From Remark 2, this implies that if a reconstruction is both *γ*- and *γ′*-parsimonious for two positive real numbers *γ* < *γ′* then it assigns 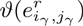 to the root.

By induction and from Corollary 1, if a *γ*-parsimonious reconstruction assigns a known value to a node *n* of *T* then it assigns known values to all nodes of the subtree rooted at *n*, except for a finite number of particular values of the asymmetry parameter *γ*.

In plain English, whatever the function *ϕ* and the parameter *γ*, the TDALP reconstructs all the unknown states with values taken among the known states, except for some degenerate cases. In particular, this holds for the Wagner parsimony.

## 4 Parametric reconstruction

### 4.1 Definition

Let us start with a remark which follows essentially from the proof of Theorem 2

**Remark 3**. *Let r be the root of T and* 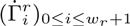 *be the elements of*

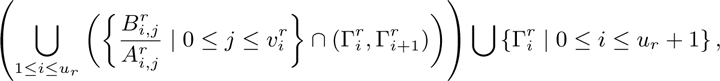

*indexed in increasing order (we have that* 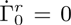 *and* 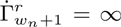). *There exists a sequence* 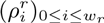 *of increasing values in ϑ*(𝒦) *such that for all* 0 ≤ *i* ≤ *w*_*r*_ *and all* 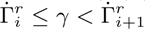, *a γ-parsimonious reconstruction associates* 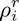 *to r. If* 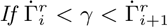 *then all γ-parsimonious reconstructions associate* 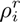 *to r*.

From Remark 3 and by induction with Corollary 1, we get that, for all unknown node *n* of T, there exists a triple 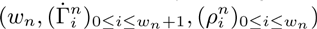 such that 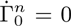, 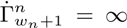, and, for all 0 ≤ *i* ≤ *w*_*n*_ and all 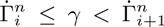, a *γ*-parsimonious reconstruction associates 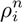 to *n*. The triple 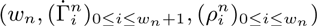 will be referred to as the *parametric reconstruction* of *n*.

A graphical, and useful, way to represent a parametric reconstruction is to cut a quarter pie at each slope 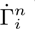 for 1 ≤ *i* ≤ *w*_*n*_ (all these slopes are in the positive quadrant). Each slice of the quarter pie is then associated to a reconstructed value for *n* and its size reflects the proportion of asymmetry parameters leading to it (Figure. 1-top-left, 2 and 3).

### 4.2 Algorithm

Theorems 1 and 2, Corollary 1 and Remark 3 suggest the approach sketched in Algorithms 1 and 2, for computing the parametric time-dependent-asymmetric-linear reconstruction of a tree *T* with regard to a known function *ϑ*. Algorithm 1 presents two functions which compute the cost and stem cost functions. These functions are used by the function main of Algorithm 2 for computing the parametric reconstruction 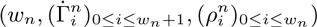 of all nodes *n* of *T*.

The complexity of the computation of the parametric reconstruction depends on the size of the piecewise linear functions (*f*_*n*_)*n∈T* and 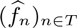, i.e. the number of intervals required to define them. Though the number of state-bounds is, by construction, smaller than |𝒦|+2, bounding the number of asymmetry-bounds is not that straightforward.

### 4.3 Bounding *u*_*n*_

From now on, we assume without loss of generality that all the bounds of the piecewise linear functions that we will consider, are necessary, in the sense that, under the notations of Theorem 1 and for all nodes *n* of *T*, the cost functions *f_n_* are such that:

- for all 0 ≤ *i* ≤ *u*_*n*_ and all 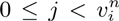, we have that 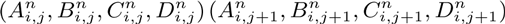;
- for all 0 ≤ *i* ≤ *u*_*n*_, there exists 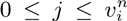 and 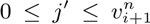 such that 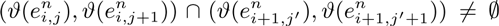 and 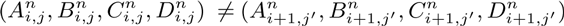;

and that the same holds for all stem cost functions 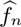. For all 0 ≤ *i* ≤ *u*_*n*_, all 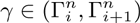 and all 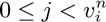, we will say that 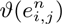 is a state-bound wrt *γ*.

For bounding the complexity of Algorithm 2, we shall bound the number of intervals in which one has to split the domain of *γ*, in order to express the map *f*_*n*_. By examining the proof of Theorem 1 and Algorithm 1, we first remark that the only stage in which the number of asymmetry-bounds increases, is that computing 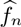 from *f*_*n*_ for the unknown internal nodes *n*. The main point of this section is thus to evaluate the maximal increase between *u*_*n*_ and 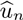. Let us start with a technical lemma.

#### Lemma 1.

*Let n be an unknown node of T. For all x* ∈ ℝ, *the x-coefficient of f*_*n*_ (*γ, x*) (*resp. of* 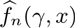) *decreases with γ*.

*Proof*. We prove the lemma by using again the standard scheme. All the base cases are plain, and so is Step 1.

Let us skip to Step 2 and assume that, for all children *m* of *n*, the *x*-coefficient of *f*_*m*_ (*γ, x*) decreases with *γ*. From the proof of Theorem 1 and for all *γ* > 0, if *i* is such that 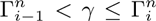, there exists an index 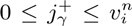 such that, by setting 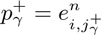, the *x*-coefficient of 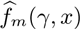 is equal to 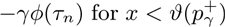 and greater than –*γϕ*(*τ*_*n*_) otherwise. Symmetrically, there exists an index 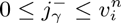 such that by setting 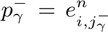, the *x*-coefficient of 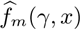 is equal to *ϕ*(*τ*_*n*_) for 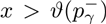 and smaller than *ϕ*(*τ*_*n*_) otherwise. The definitions of these nodes altogether with the induction hypothesis implies that the values 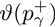 and 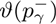 both increase with *γ*.

Let now consider two positive real numbers 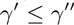. We then have 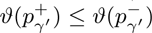, 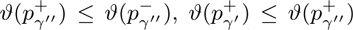 and 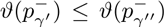. The first two inequalities come from construction (proof of Theorem 1) and the two last ones from the induction hypothesis. This leaves only two cases to investigate:

1. 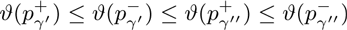 and
2. 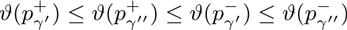.

With regard to the value of *x*, five possibilities arise in Case 1:

- if 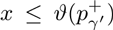 then the *x*-coefficients of 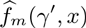 and 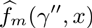 are equal to 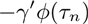 and 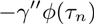 respectively;
- if 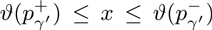 then the *x*-coefficient of 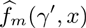 is equal to that of *f*_*m*_(*γ′,x*), thus greater than *γ′ϕ*(*τ*_*n*_), and the *x*-coefficient of 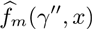 is equal to *–γ″ϕ*(*τ*_*n*_);
- if 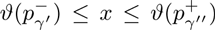 then the*x*-coefficients of 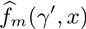 and 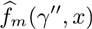 are *ϕ*(*τ*_*n*_) and *–γ″ϕ*(*τ*_*n*_) respectively;
- if 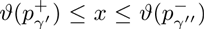 then the *x*-coefficient of 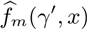 is equal to *ϕ*(*τ*_*n*_) and that of 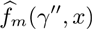 is equal to that of *f*_*m*_(*γ″,x*), thus smaller than *ϕ*(*τ*_*n*_);
- if 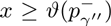 then the *x*-coefficients of 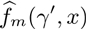 and 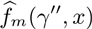 are both equal to *ϕ*(*τ*_*n*_).

Similarly, five possibilities arise in Case 2:

- if 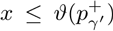 then the *x*-coefficients of 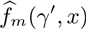 and 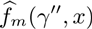 are equal to *–γ′ϕ*(*τ*_*n*_) and *–γ″ϕ*(*τ*_*n*_) respectively;
- if 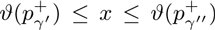 then the *x*-coefficient of 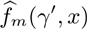 is equal to that of *f*_*m*_(*γ′, x*), thus greater than *–γ′ϕ*(*τ*_*n*_), and the *x*-coefficient of 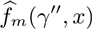 is equal to *–γ″ϕ*(*τ*_*n*_);
- if 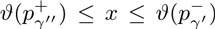then the*x*-coefficients of 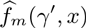and 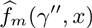 are equal to that of *f*_*m*_(*γ′,x*) and *f*_*m*_*(γ″,x)*, respectively;
- if 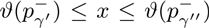 then the *x*-coefficient of 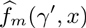 is *ϕ*(*τ*_*n*_) and that of 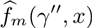 is equal to that of *f*_*m*_(*γ″,x*), thus smaller than *ϕ*(*τ*_*n*_);
- if 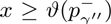 then the *x*-coefficients of 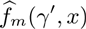 and 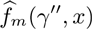 are both equal to *ϕ*(*τ*_*n*_).

In all the situations cover by Cases 1 and 2, the *x*-coefficient of 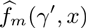 is always greater than that of 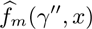, either directly or from the induction hypothesis. The *x*-coefficient of 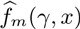 does decrease with *γ*.

Step 3 follows straightforwardly from Equation 2 and yields to conclude that, for all reals *x*, the *x*-coefficient of *f*_*n*_(*γ,x*) decreases with *γ*.

#### Algorithm 1

Computation of *f*_*n*_ and 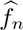 for all nodes *n* of *T* (under the notations of Theorem 1).

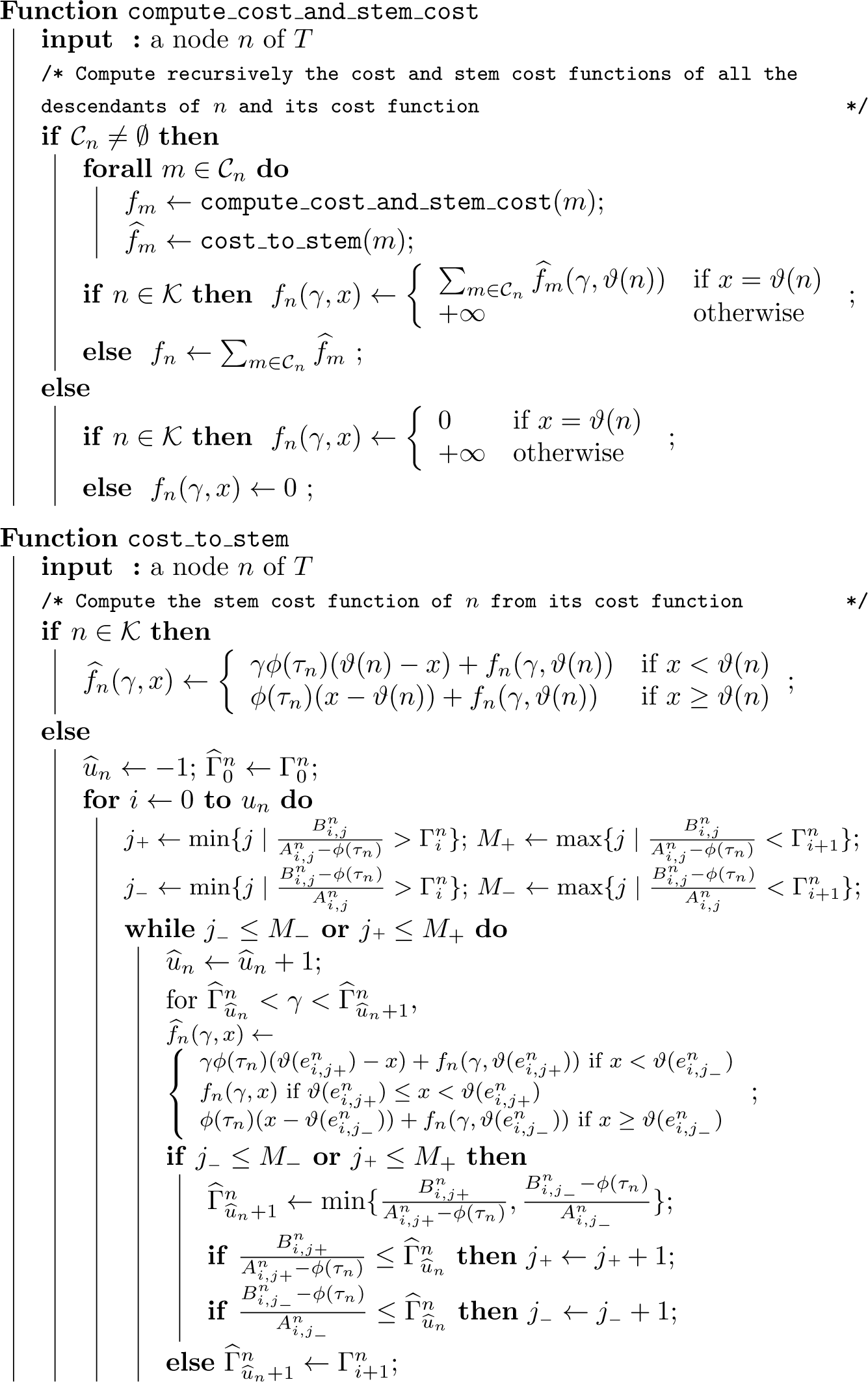

#### Algorithm 2

Parametric reconstruction of a tree *T*.

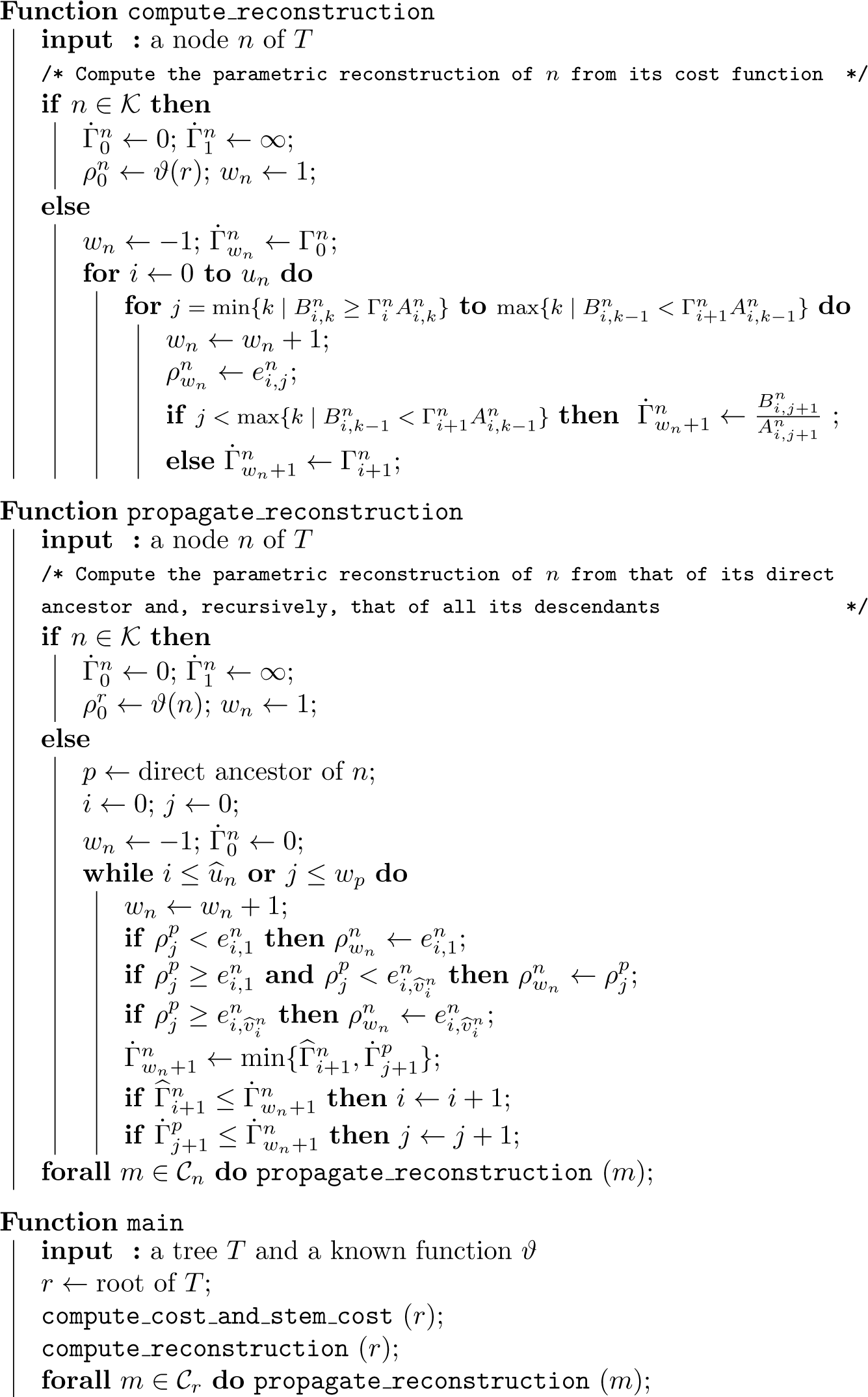

**Figure 1:**
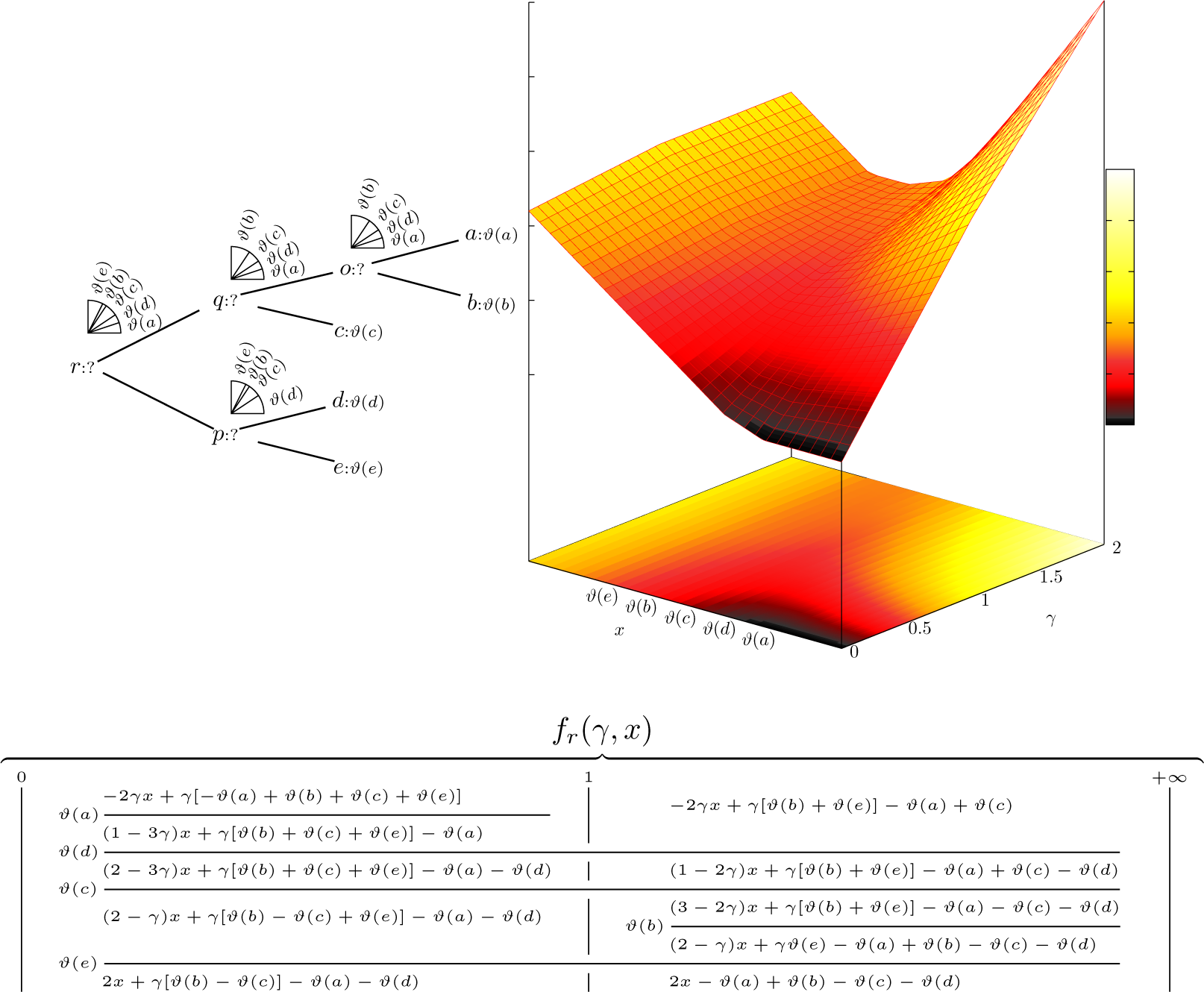
Top: A tree in which only leaves are known with *ϑ*(*a*) < *ϑ*(*d*) < *ϑ*(*c*) < *ϑ*(*b*) < *ϑ*(*e*) and where the pie representations of the reconstructed values according to the the slope *γ* are above the unknown nodes (Left) Graphic representation of the subtree cost function of its root (Right). Bottom: The subtree cost function of its root. The subtree cost function is that of the generalized parsimony scheme of [1] (i.e. with *ϕ*(*τ*) = 1).

#### Lemma 2.

*Let n be a non-root unknown internal node n of T with* 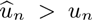, *l be an index in* 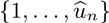 *such that* 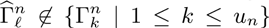 *and I be such that* 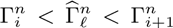. *There exists an index* 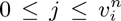 *such that at least one of the following assertions holds*:

1. 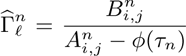 *and* 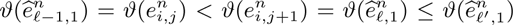 *for all* 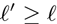;
2. 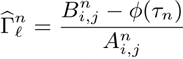 *and* 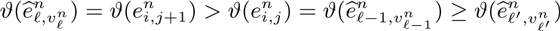 *for all* 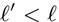.

*Proof.* From the proof of Theorem 1 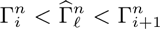 then there exists an index *j* such that 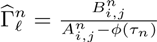 or 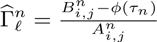.

If 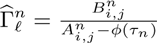 and still from the proof of Theorem 1, we have that 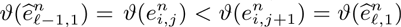. Lemma 1 then ensures that for all *i′* ≥ *i* and all *j'* such that 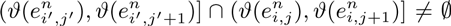, we have that 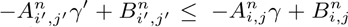 and all 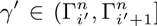 and all 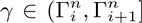. It follows that, if 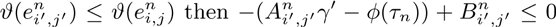 for all 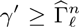. This implies that 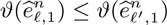 for all 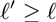.

The situation is symmetrical if 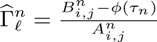.

In plain English, Lemma 2 says that each time that an asymmetry-bound 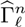 is created, either 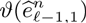 is no longer required as state-bound for all 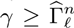, (i.e. 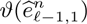 vanishes after 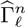), or 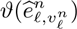 was not a necessary state-bound for all 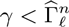 (i.e. 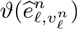appears after 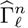). Since all the state-bounds of *f*_*n*_ belong to *ϑ*(𝒦_*n*_) and, from Lemma 2, appear and vanish at most once during the computation of 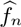 from *f*_*n*_, we have that

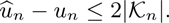

#### Theorem 3.

*For all non-root nodes n of T, we have that*

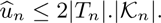

*Proof.* Each asymmetry-bound Λ of 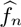 is such that there exists an unknown node *q* ϵ *T*_*n*_, such that Λ is an asymmetry-bound of 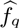 but not of *f*_*q*_. From Lemma 2, this implies that there exists a known node *k* in *ϑ*(𝒦_*q*_) such that *ϑ*(*k*) is required or not as state-bound of 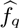 according to whether the asymmetry-parameter is smaller to Λ or not (or conversely). In short, all asymmetry-bounds of 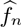 involve an unknown node *q* ϵ *T*_*n*_ and a known descendant *k* of *q*. Moreover, from Lemma 2, all known descendants *k* of a known node *q* can be involved with at most twice asymmetry-bounds of 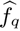 which are not asymmetry-bounds of *f*_*q*_. Since each node of 𝒦_*n*_ has less than |*T*_*n*_| unknown ancestors in *T*_*n*_, it follows that the total number of asymmetry-bounds of 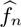 cannot exceed 2|*Tn*|.|𝒦_*n*_|.

*Corollary 2.*Under the assumption that the number of children of node is bounded independently of the size of the tree, the algorithmic complexity of the computation of the time-dependent-asymmetric-linear parametric reconstruction of a tree T with a set of known nodes 𝒦 is O*(|*T*|*^2^*.|*𝒦*|*^2^) both in time and memory space.

*Proof.* For all non-root unknown nodes *n* of *T*, computing the stem cost function 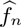 from the cost function *f*_*n*_ is linear with the total number of “pieces” required to define 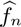 (i.e. its size, see Algorithm 1). From Theorem 3, less than 2|*T*_*n*_|.|𝒦_n_| asymmetry-bounds are required for *f*_*n*_. Over all intervals between two successive asymmetry-bounds, there are at most |𝒦_*n*_| state-bounds required. It follows that the size of 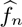 is smaller than 2|*T*_*n*_|.|𝒦_*n*_|^2^ and its computation from *f*_*n*_ is *O*(|*T*_*n*_|.|𝒦_*n*_|^2^).

Obtaining the cost function *f_n_*for all the stem cost functions 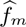 of its children *m* is performed by using a procedure similar to that merging sorted lists. Under the assumption that the number of children *m* is bounded independently of the size of the tree, this operation is linear with total size of the stem cost functions 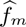. In short, this stage is *O*(|*T*_*n*_|.|𝒦_*n*_|^2^).

It follows that the computation of all the cost and stem cost functions of the unknown nodes of *T* is *O*(|*T*|^2^.|𝒦|^2^) both in time and memory space.

The stage computing the parametric reconstructions from the cost and stem cost functions is linear with their total size (Algorithm 2), which gives us the overall complexity of the algorithm.

### 5 Example

The TDALP was applied to the dataset of [10], which contains the phylogenetic tree of extant cetaceans (including branch lengths) and their (average) body sizes (this dataset was used as it was - we just pruned extant taxa of which the body size was not provided). The results of the parametric reconstruction of the body size of cetaceans are displayed in Figure. 2 and 3. Figure 2 shows the whole phylogenetic tree with the quarter pie representation of the parametric reconstruction of all the nodes. Figure 3 details the parametric reconstruction of the most recent common ancestor of cetaceans (i.e. the root of the tree of Figure 2).

**Figure 2:**
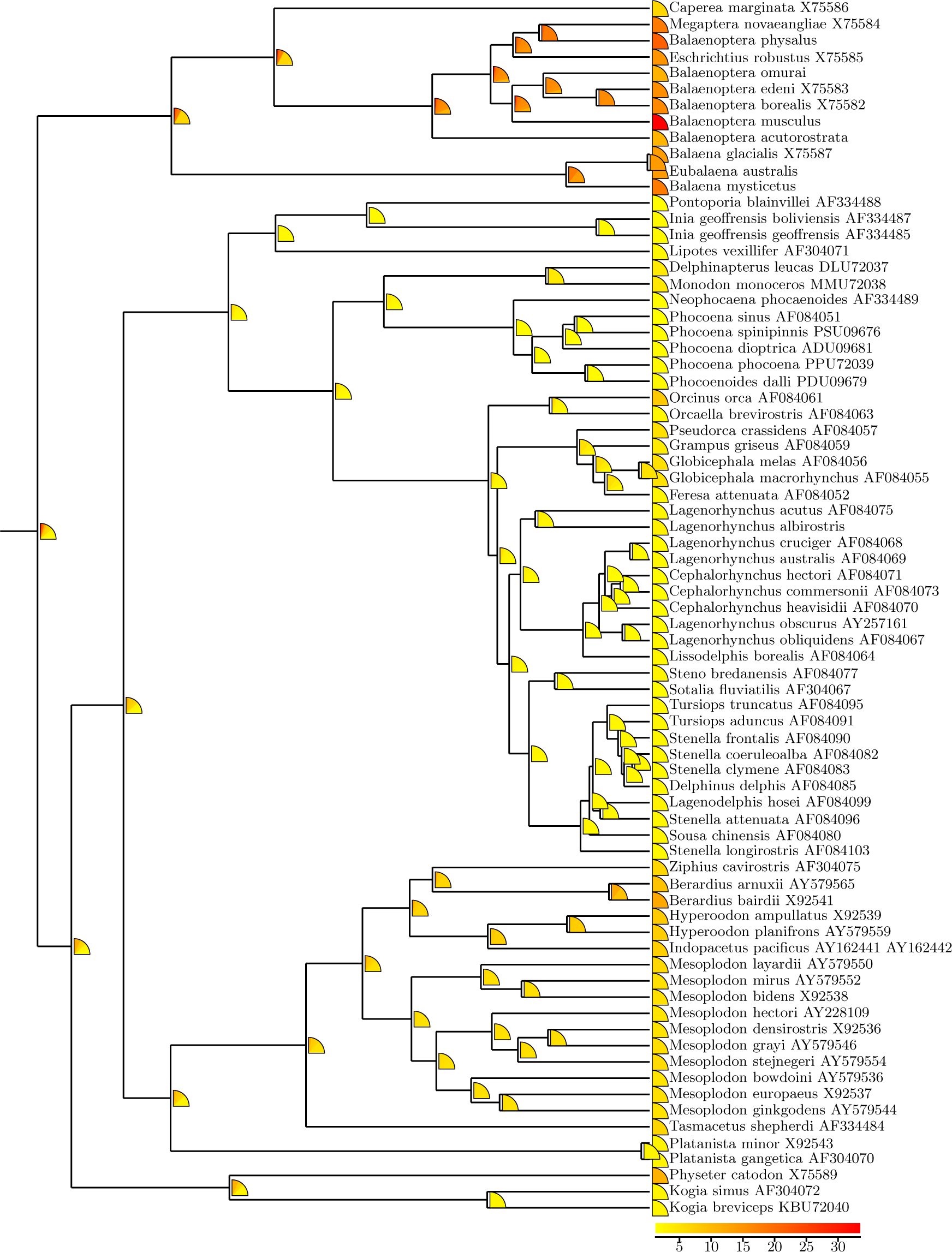
Parametric reconstruction of cetacean body size in meters, by taking into account branch lengths with 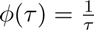. The reconstructed states are represented by colors in the quarter pies figuring the parametric reconstruction. The color scale is displayed at the bottom-right of the figure.

**Figure 3:**
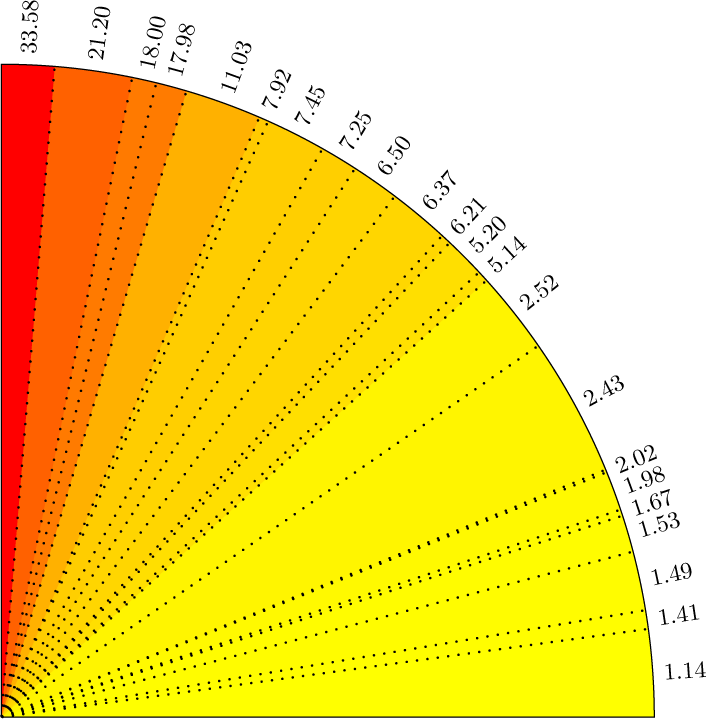
Detail of the parametric reconstruction of the root of the cetacean phylogenetic tree displayed in Figure 2.

Figure 2 provides a synthetic representation of all the possible parsimonious reconstructions of the ancestral states. One can see at a glance that is little uncertainty with the reconstruction of some of the nodes while that of other ones is more mixed. This representation actually contains all the possible reconstructions from the TDALP with regard to the asymmetry parameter.

By focusing on a node of interest, for instance the root of the tree as displayed in Figure 3, one can take a closer look on which reconstructed states are possible, in what extent they are supported, and which are the corresponding evolutionary assumptions. In Figure 3, we do observe that a root state smaller than 2.52 m is supported by approximately half of the asymmetry parameters, which corresponds to trends ranging from highly positive to neutral. Conversely, an ancestral state greater than 8 m is reconstructed only for the uppermost quarter of the asymmetry parameters, which corresponds to the most negative evolutionary trends.

Remark that any assumption, or any evidence (e.g. some fossils), about one or several ancestral states directly translates into an assumption or an evidence about the asymmetry parameter, thus in a certain sense, about the nature and the intensity of the underlying evolutionary trend. This is basically done by checking which interval of the corresponding parametric reconstruction contains the assumed or known ancestral state.

